# The role of sphingosine-1-phosphate receptor 2 in mouse retina light responses

**DOI:** 10.1101/2023.09.01.555709

**Authors:** Abhishek P Shrestha, Megan Stiles, Richard C. Grambergs, Johane M. Boff, Saivikram Madireddy, Koushik Mondal, Rhea Rajmanna, Hunter Porter, David Sherry, Richard L. Proia, Thirumalini Vaithianathan, Nawajes Mandal

## Abstract

The bioactive sphingolipid sphingosine-1-phosphate (S1P) acts as a ligand for a family of G protein-coupled S1P receptors (S1PR1-5) to participate in a variety of signaling pathways. However, their specific roles in the neural retina remain unclear. We previously showed that S1P receptor subtype 2 (S1PR2) is expressed in murine retinas, primarily in photoreceptors and bipolar cells, and its expression is altered by retinal stress. This study aims to elucidate the role of S1PR2 in the mouse retina. We examined light responses by electroretinography (ERG), structural differences by optical coherence tomography (OCT), and protein levels by immunohistochemistry (IHC) in wild-type (WT) and S1PR2 knockout (KO) mice at various ages between 3 and 6 months. We found that a- and b-wave responses significantly increased at flash intensities between 400∼2000 and 4∼2,000 cd.s/m^2^ respectively, in S1PR2 KO mice relative to those of WT controls at baseline. S1PR2 KO mice also exhibited significantly increased retinal nerve fiber layer (RNFL) and outer plexiform layer (OPL) thickness by OCT relative to the WT. Finally, in S1PR2 KO mice, we observed differential labeling of synaptic markers by immunohistochemistry (IHC) and quantitative reverse transcription polymerase chain reaction (RT-qPCR). These results suggest a specific involvement of S1PR2 in the structure and synaptic organization of the retina and a potential role in light-mediated functioning of the retina.

## Introduction

Neuronal membranes are enriched in sphingolipids,[1] which serve pivotal roles in structure and signaling. The bioactive lipid sphingosine-1-phosphate (S1P) is a sphingosine metabolite produced by two sphingosine kinase isoenzymes.[2] S1P plays important roles in synaptic transmission, modulation of neurotransmitter release, membrane excitability, and synaptic strength in the central nervous system. Although sphingolipids are possible mediators of retinal diseases, including diabetic retinopathy[3] and age-related macular degeneration[4] and of retinal degenerative processes in general, [5] their roles in retinal function are unclear.

The bioactive potential of S1P is primarily derived from its interactions with a family of G protein-coupled S1P receptors (S1PR1-S1PR5) that are involved in signaling via numerous signaling pathways.[6] Their expression patterns differ by tissue localization and cell specialization.[7, 8] We previously demonstrated the expression of *S1PR2* mRNA in the murine neural retina, albeit at lower expression levels than observed for other S1P receptors. In rat retinas, we observed expression of S1PR2 protein primarily within photoreceptor (PR) inner segments and the inner nuclear layer (INL) by immunohistochemistry (IHC).[1] The expression variability of S1PR2 in different regions of the murine retina could suggest the involvement of S1PR2 signaling in aspects of PR and bipolar cell function, but the mechanism by which S1PR2 affects retinal function remains unknown.

The purpose of this study was to determine the role of S1PR2 in the retina at the functional and cellular level using S1PR2 knockout mice (S1PR2 KO). We investigated the functional and structural integrity of the retina in WT and S1PR2 KO mice using electroretinography (ERG) analysis and optical coherence tomography (OCT). Further, we also compared the expression of synaptic genes and proteins in WT and S1PR2 KO mouse retinas by immunohistochemistry (IHC) and quantitative reverse transcription polymerase chain reaction (RT-qPCR). We observed the specific involvement of S1PR2 in the structure and synaptic organization of the retina and its potential role in light-mediated retinal function.

## Materials and Methods

### Mice

All methodological procedures were performed in agreement with the Association for Research in Vision and Ophthalmology (ARVO) Statement for the use of Animals in Ophthalmic and Vision Research and the University of Tennessee Health Science Center (UTHSC) Guidelines for Animals in Research. All procedures, euthanasia (CO2 asphyxiation), and tissue extraction methods were reviewed and approved by the UTHSC Institutional Animal Care and Use Committee (IACUC; Approval # 19-0104). Albino S1PR2 KO and WT littermate male and female mice were raised under cyclic dim-light (5-20 lux) and dark conditions (12/12 h). Pigmented S1PR2 KO mice were obtained from Dr. Richard Proia (NIDDK, NIH) and an albino line was generated by a minimum of seven generations of backcrossing with Balb/c mice.[9] PCR genotyping was used to confirm the genotypes of the original and albino lines. The mice underwent assessments of retinal structure and function, including OCT, IHC for neuronal and synaptic markers, RT-qPCR assays for expression of retinal cell markers and sphingolipid metabolic genes, and ERG measurements at various ages between 3 and 6 months.

### Electroretinography (ERG)

Visual function of WT and S1PR2 KO mice at 3 and 6 months of age was measured by light- and dark-adapted ERG. Scotopic and photopic flash ERGs were recorded with a Diagnosys Espion E2 ERG system (Diagnosys LLC, Lowell, MA) as previously described [10] flash ERGs were recorded for both eyes. Scotopic ERG was used to assess rod photoreceptor function, with five strobe flash stimuli presented at intensities of 0.0004, 0.04, 4, 400, and 2000 candela (cd.s/m^2^). The a-wave reflects the physiological health of the photoreceptors in the outer retina, while the b-wave reflects the health of the inner retinal layers and, thus, the activity of ON bipolar cells that respond to the onset of light.[11, 12] The amplitude of the a-wave was measured from the pre-stimulus baseline to the a-wave trough. The scotopic b-wave response was measured for secondary neurons where the amplitude of the b-wave was measured from the trough of the a-wave to the peak of the b-wave. For the evaluation of cone photoreceptor function (photopic ERG), two strobe flash stimuli at 3.0 and 10.0 cd.s/m^2^ were presented to dilated, light-adapted (5 minutes at 2 cd.s/m^2^) mice followed by flickers at 10 Hz under a steady adapting field of 2 cd.s/m^2^.

### SD-OCT analysis of retinal layers in mice

The thickness of different retinal layers was measured using spectral-domain optical coherence tomography (SD-OCT) with a Bioptigen Envisu Image Guided SD-OCT system (Lecia Microsystems, Buffalo Grove, IL). Briefly, 3 and 6-month-old mice from both genotypes (WT and S1PR2 KO) were anesthetized and placed on the imaging platform under room light. We performed radial line scans of 2-mm centered at the optic nerve head. Horizontal and vertical SD-OCT images from the line scans were exported in TIF format. Automated image segmentation software provided by the manufacturer was used for the segmentation of the images, and each segmentation was manually verified. Thickness data was collected from an equidistance region between 0.6-0.8 mm away from the center of the optic nerve on both sides of the image. Therefore, four readings were collected from the two images (horizontal and vertical) per eye. Using GraphPad Prism software the means were compared using unpaired t test with Welch’s correction. Statistical significance was accepted at P < 0.05.

### Immunohistochemistry (IHC)

Differences in the expression and localization of neuronal and synaptic markers in harvested retinas were visualized by IHC. Fixed eyes were thoroughly washed in phosphate-buffered saline (PBS), cryoprotected in PBS with three increasing concentrations of sucrose (10%, 20%, and 30%), embedded in optimal cutting temperature (OCT) embedding medium (Fisher Healthcare, Houston, TX), placed in a Tissue-Tek Cryomold (Fisher Scientific, Pittsburg, PA), snap frozen, and stored at -80°C. Sections of 15 μm thickness were cut in a Leica CM1850 cryostat (Leica Microsystems, Bannockburn, IL, USA) with a chamber temperature of −12 to −15°C and the sections were placed on Superfrost™ Plus microscope slides (Fisher Scientific) and stored at a temperature of −20 to −80°C.

The sections mounted on glass slides were allowed to dry at room temperature for 2 hours. All the immunoassay was performed from sections obtained in the central retina and kept consistent between groups. A PAP pen liquid blocker (Electron Microscopy Sciences, Hatfield, PA) was applied at the edges of the slides to generate a hydrophobic barrier that was air-dried for 15 minutes. The prepared slides were washed three times with PBS, permeabilized, blocked with Perm blocking solution (0.3% Triton-X-100 and 5% donkey serum in PBS), and incubated for 2 hours at 4°C in a hydrated chamber. Primary antibodies (**Table 1**) were diluted in blocking solution and applied to the slides, which were incubated overnight at 4°C; primary antibodies were omitted for the negative controls. After washing with PBS, secondary antibodies conjugated to the indicated fluorophores (**Table 2**) were applied (1:500 dilutions) and incubated for 2 hours at room temperature (RT). For negative controls, primary antibodies were omitted. The slides were washed once in PBS-Tween, thrice in PBS, and once in distilled water, and coverslips were attached using Prolong Diamond Antifade Mountant (ThermoFisher Scientific, Waltham, MA).

**Table 1.** Primary antibodies used for IHC.

| **Antigen** | **Antiserum** | **Host** | **Dilution** | **Source (Number)** | **Marker for** |
| --- | --- | --- | --- | --- | --- |
| Rhodopsin | 4D2,<br>monoclonal | Mouse | 1:500 | Sigma (MABN15) | Rod photoreceptors |
| Cone Arrestin | Polyclonal<br>anti- Cone<br>Arrestin<br>n | Rabbit | 1:1000 | Sigma (AB15282MI) | Cone photoreceptors |
| Secretagogin | Polyclonal<br>anti-<br>secretagogin | Sheep | 1:100 | Fisher (50-165-6430) | Cone-bipolar cells |
| PKC | Monoclonal<br>anti-PKC | Mouse | 1:250 | Santa Cruz<br>(sc-17769) | Rod-type bipolar cells |
| Ribeye | Polyclonal<br>anti-ribeye | Guinea<br>pig | 1:500 | Synaptic Systems,<br>#192 104 | Synaptic ribbons (IPL) |
| Calbindin | Monoclonal<br>anti-calbindin | Mouse | 1:500 | ThermoFisher<br>(MA5-32993) | Horizontal cells |
| Pan-MAGUK | Monoclonal<br>pan-MAGUK,<br>clone K28/86 | Mouse | 1:250 | Millipore Sigma<br>(MABN72MI) | Postsynaptic excitatory<br>glutamatergic synapses |
| Synaptophysin | Polyclonal<br>anti-synaptophysin | Rabbit | 1:250 | Alomone lab<br>(ANR-013) | Synaptic vesicles |
| SV2 | Monoclonal<br>anti-SV2 | Mouse | 1:250 | DSHB, #SV2 | Synaptic vesicles |

**Table 2.** Secondary antibodies used for IHC.

| **Antibody** | **Conjugation** | **Source (Number)** |
| --- | --- | --- |
| Donkey anti-guinea pig | Cy3 | Jackson ImmunoResearch, #706-165-148 |
| Donkey anti-mouse | Alexa Fluor 647 | Southern Biotech, #6440-31 |
| Donkey anti-rabbit | Alexa Fluor 488 | Southern Biotech, #6440-30 |
| Donkey Anti-Sheep | Alexa Fluor 488 | ThermoFisher, A11015 |

IHC images were obtained using the 60 X silicon objective with 1-7 x software zoom and 0.45 μm z-step size on an Olympus FV-3000 laser-scanning confocal system (Olympus, Shinjuku, Tokyo, Japan) or an Olympus IX-83 inverted microscope controlled by Olympus FV31S-SW software (Version 2.3.1.163) with a Galvano scanner. For each image, we obtained three sequential line scans at 10.0 μs/pixel with a scan size of 256 × 256 pixels. Analysis of the resulting images was performed with ImageJ software (NIH) with constant acquisition parameters for pinhole diameter, laser power, PMT gain, scan speed, optical zoom, offset, and step size.

#### Area of expression

We analyzed three images from each animal with Image J software to obtain the percentage area labeled by the antibody. Images were first split into RGB channels, a common threshold in all images was set for each channel, and a defined region of interest was used to obtain the percent area of pixels over the threshold. Values per retina were averaged and plotted as mean ± SEM (standard error of the mean).

#### Cell Counting

Cell soma were labeled for 5 min with 300 nM DAPI, slides were washed once in PBS-Tween, three times in PBS, and once in distilled water), and coverslips were mounted with Prolong Diamond Antifade Mountant (ThermoFisher Scientific). The round shape of the ONL, INL, or GCL cell nucleus was considered to be a neuron in the respective nuclear layer. Digital images corresponding to different retinal regions were overlaid with random boxes 100 × 100 mm in size, and the number of cells in each layer was counted using the cell counter plugin of the Fiji/ImageJ software.

### Quantitative reverse transcription polymerase chain reaction (RT-qPCR)

Differences in retinal gene expression were determined by RT-qPCR of harvested retinas. Briefly, S1PR2 KO and WT mice were euthanized under room light, retinal tissue was isolated and frozen, and total cellular RNA was isolated from thawed tissue using a PureLink^®^ Micro-to Midi Total RNA Purification System (Invitrogen, Carlsbad, CA) following the manufacturer’s protocol. RNA was quantitated and equal amounts (1.0 µg) of RNA from each sample were converted to first-strand cDNA using SuperScript III First-Strand Synthesis SuperMix (Invitrogen, Carlsbad, CA, USA). This first-strand cDNA was used for RT-qPCR analysis with intron-spanning primers that had been generated using Primer 3 software and custom synthesized. Their sequences are shown in **Table 3**. RT-qPCR and melt-curve analyses were performed using iQ SYBR Green Supermix (Bio-Rad, Hercules, CA) and a Bio-Rad iCycler optical system. Expression of genes of interest relative to the expression of housekeeping genes in different samples was quantitated using the comparative Ct (threshold cycle) value method.[1, 10, 13–15]

**Table 3.**
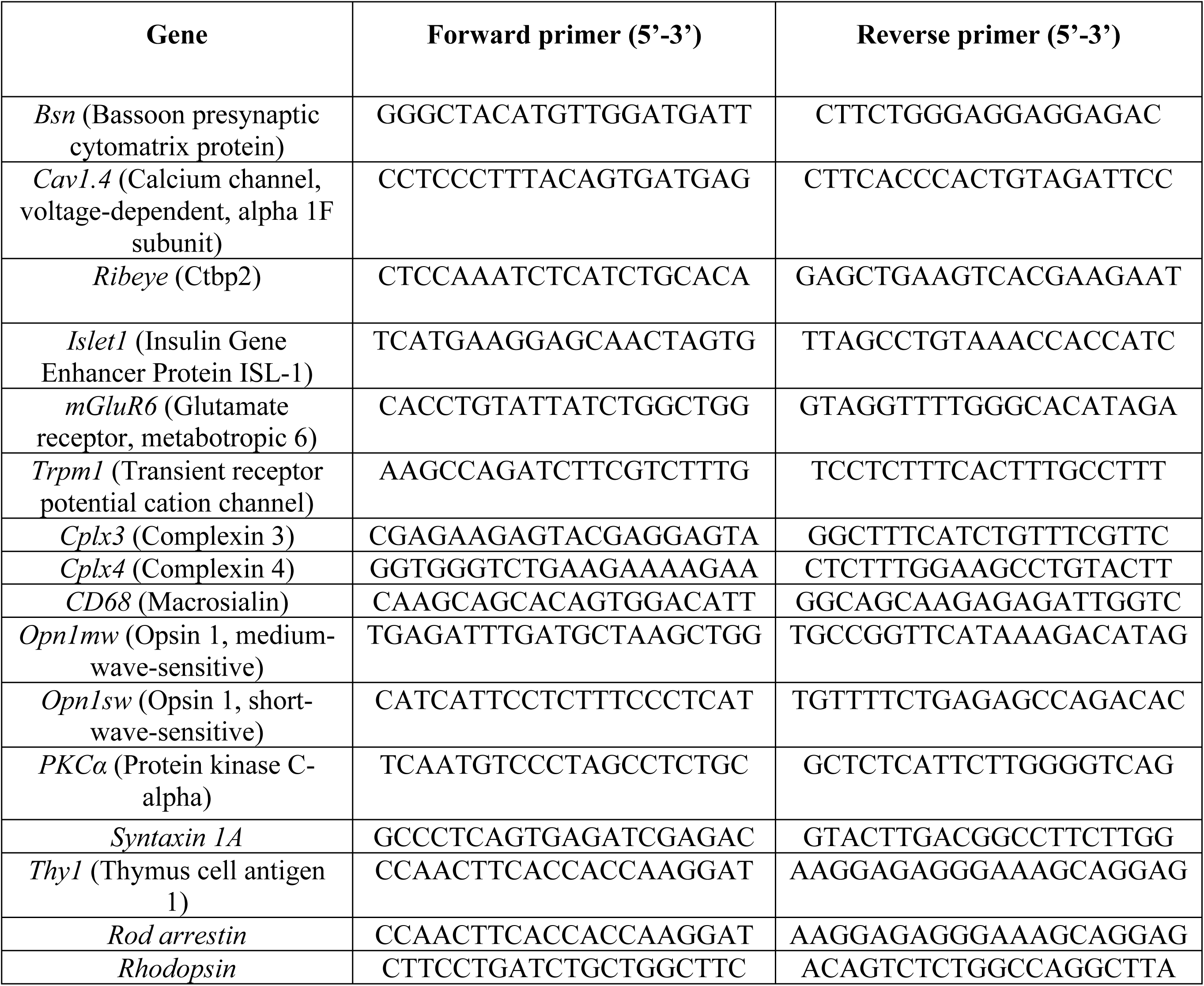
Sequence of primers designed to measure the gene expression by qRT-PCR from mouse retina.

## Results

### Role of S1PR2 in light responses of the retina

We first examined the functional integrity of the retina in WT and S1PR2 KO mice using full-field ERG (ffERG). Because ffERG wave components and underlying sources are governed by different states of adaptation and the strength of the stimulus flash intensity, we compared scotopic (**Figures 1A-C**) and photopic (**Figures 1D-F**) ERG measurements to determine the specific roles of S1PR2 in rod vs. cone-pathway function. In dark-adapted retinas at lower flash intensities (<4 cd.s/m^2^), we found no differences in the a-wave amplitude of the baseline ERG responses of WT and S1PR2 KO mice. However, at higher flash intensities (400 and 2000 cd.s/m^2^), S1PR2 KO mice displayed significantly higher scotopic a-wave amplitude than did WT mice at both three (**Supplement Figure S1A**) six months of age (**Figure 1B**) and their scotopic b-wave ERG responses were also significantly higher than those of WT mice at even lower flash intensities than examined for the a-wave responses (4, 400, and 2000 cd.s/m^2^), as shown in **Supplement Figure S1B** and **Figure 2C**. We also observed that the retina from light-adapted S1PR2 KO mice exhibited a significantly augmented amplitude of photopic b-wave responses relative to that of the WT mice (**Figure 1F**). These results underscore the importance of S1PR2 in light responses and synaptic transmission between photoreceptor-bipolar cells, and no difference was noticed between 3 and 6 months of age.

**Figure 1.**
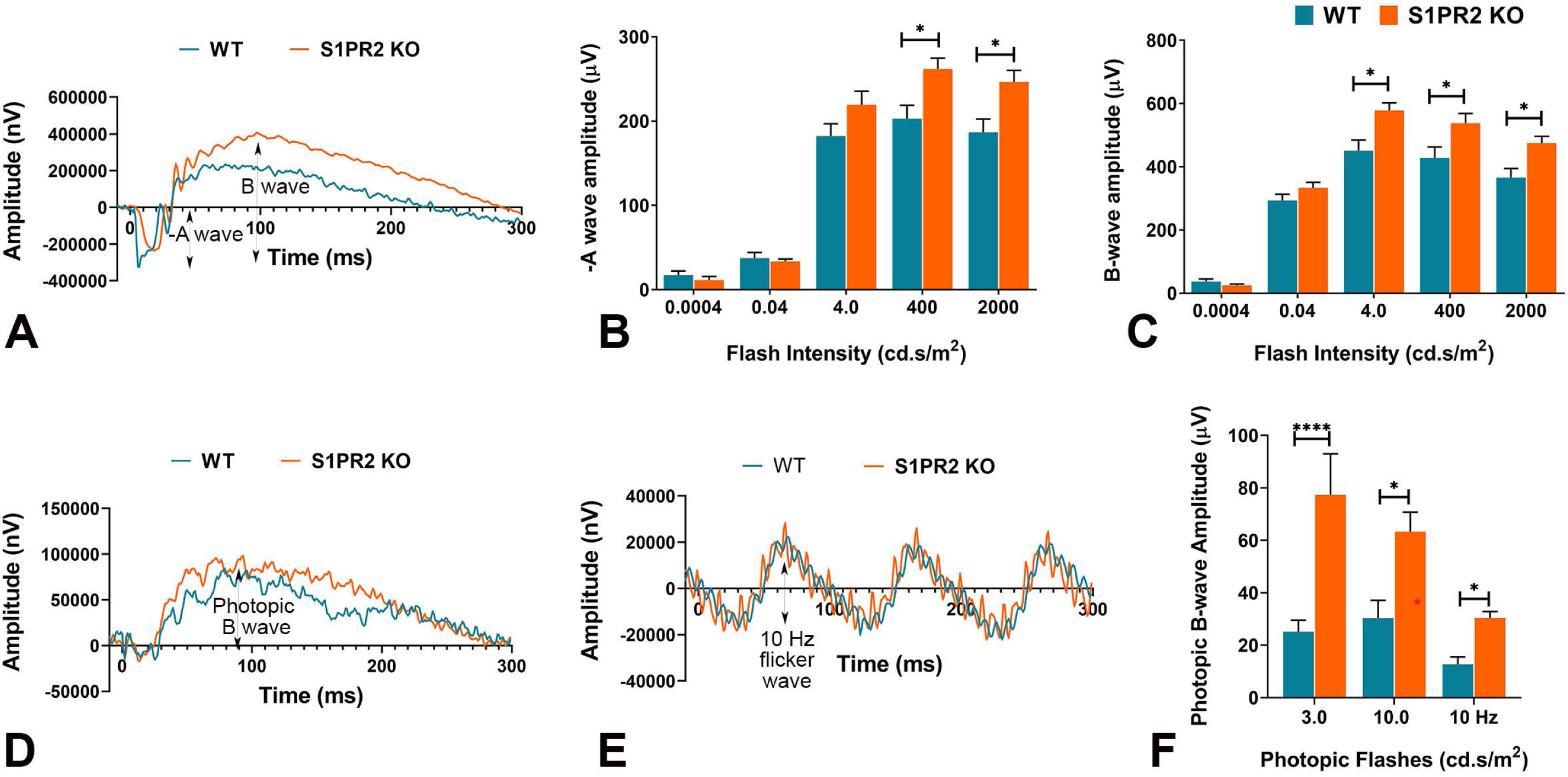
S1PR2 knockout (KO) mice exhibited higher baseline ERG responses than did wild-type (WT) mice at 6-months of age. Shown are statistical analyses of scotopic ERG A- and B-wave amplitudes and photopic B-wave at different flash intensities for 6-month-old WT (blue) and S1PR2 KO (orange) mice. **(A)** Scotopic ERG amplitude waveforms (0 to 300 ms) of WT and S1PR2 KO mice. **(B)** At higher flash intensities (400 and 2000 cd.s/m^2^), S1PR2 KO mice demonstrated significantly higher mean scotopic A-wave ERG responses than WT mice. **(C)** S1PR2 KO mice demonstrated significantly higher mean scotopic B-wave ERG responses than WT mice at 4.0, 400, and 2000 cd.s/m^2^ flash intensities. **(D)** Photopic ERG amplitude waveforms (0 to 300 ms) of WT and S1PR2 KO mice. **(E)** ERG response amplitude waveforms elicited at 10 Hz flash flicker stimulus in WT and S1PR2 KO mice. **(F)** S1PR2 KO mice displayed significantly higher mean photopic B-wave ERG response amplitudes than WT mice at 3.0, cd.s/m^2^, 10.0 cd.s/m^2^, and 10 Hz flash flickering stimulus. Shown are mean values ± SEM. \**p* < 0.05, ****P < 0.0001 indicate significant differences between groups of WT and S1PR2 KO mice (N = 8 mice per genotype). Abbreviations used: ERG, electroretinography; KO, knockout; WT, wildtype.

**Figure 2.**
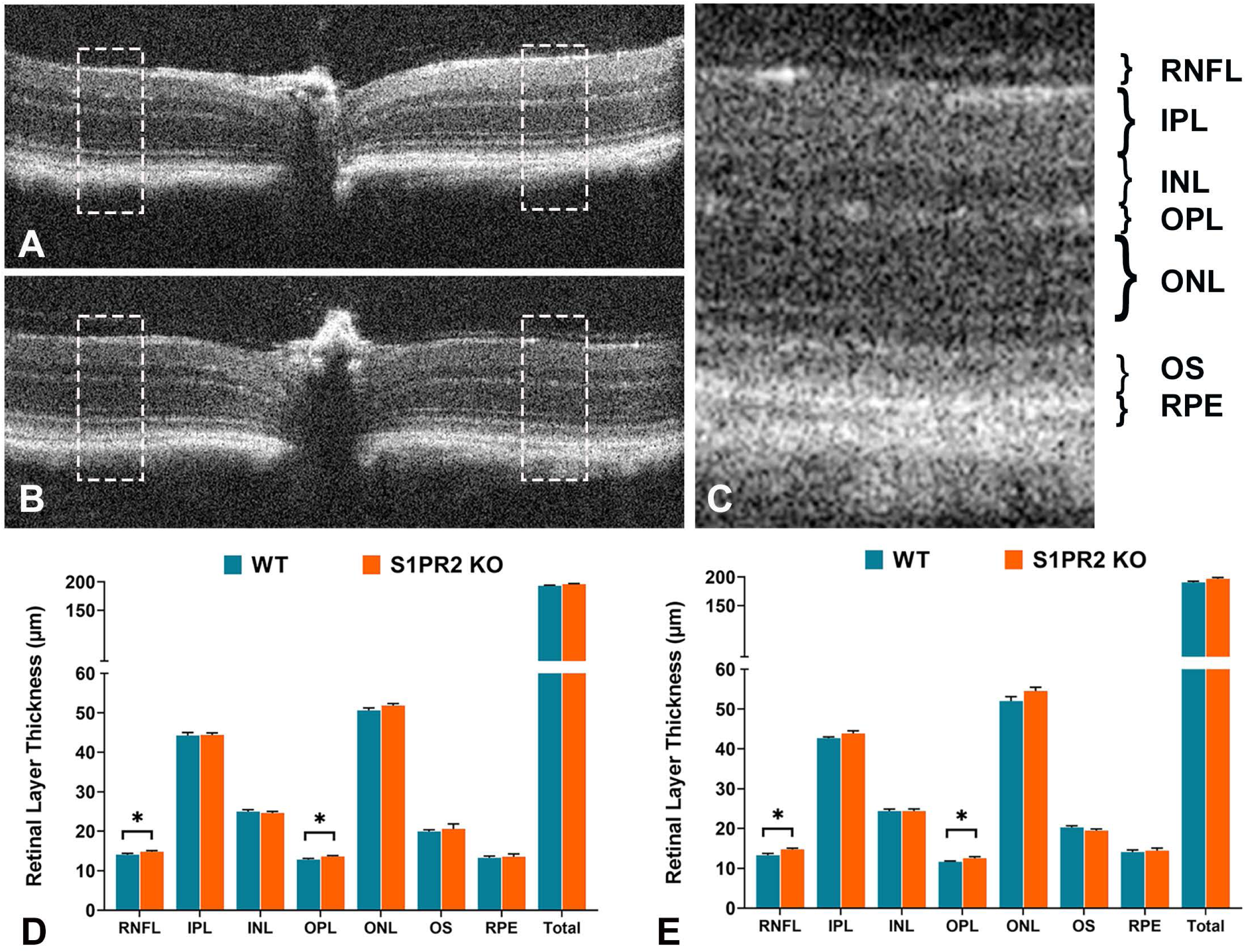
The retinas of S1PR2 KO mice exhibited increased baseline RNFL and OPL thickness relative to those of WT mice. Retinal layer thickness was measured by OCT at 3 and 6 months of age. **(A)** Representative image of a 2-mm radial scan of central retina from a WT mice and **(B)** is a representative image from S1PR2 KO mice. The boxed regions indicate the places where the thickness measures were captured. **(C)** A representative enlarged SD-OCT image of the boxed region showing the layers of the retina. (D) and (E) are thickness layers of the retina in micrometer, WT (blue) and S1PR2 KO (orange) mice, at 3 months and 6 months of age, respectively. S1PR2 KO mice exhibited significantly increased RNFL and OPL thickness relative to WT mice, while no significant differences were observed in IPL, INL, ONL, OS, RPE, and total retinal thickness. Shown are mean values ± SEM. \**p* ≤ 0.05 indicate significant difference between groups of WT and S1PR2 KO mice (N = 64; 4 reading per eye, 16 eyes from 8 mice). Abbreviations used: KO, knockout; INL, inner nuclear layer; IPL, inner plexiform layer; ONL, outer nuclear layer; OPL, outer plexiform layer; OS, outer segment; RNFL, retinal nerve fiber layer; RPE, retinal pigment epithelium; total, total thickness of all layers; WT, wildtype.

### Role of S1PR2 in the structural integrity of the retina

To reveal any morphological entities that can accommodate these altered a- and b-waves, we compared the retinal layers of age-matched WT and S1PR2 KO animals mice at both 3 months and 6 months of age using Optical Coherence Tomography (OCT). We measured the thickness of 7 segmented layers of the central retina of each eye and the total thickness at equidistance place at 0.6-0.8 mm away from the center of the optic nerve head (**Figure 2 A-C**). We found that at 3 months of age, the thickness of the retinal nerve fiber layer (RNFL) (14.83 ± 0.25 vs. 14.07 ± 0.34; p = 0.04, n = 64 [4 reading per eye, 16 eyes from 8 mice]) and outer plexiform layer (OPL) (13.62 ± 0.22 vs. 12.81 ± 0.31; p = 0.02, n = 64) in S1PR2 KO mice were significantly higher than those in WT mice (**Figure 2D**). However, we observed no changes in the inner plexiform layer (IPL), outer nuclear layer (ONL), outer segment (OS), or retinal pigment epithelium (RPE), as shown in **Figure 2D**. We observed the similar changes at 6 months old mice; RNFL (14.76 ± 0.30 vs. 13.28 ± 0.47; p = 0.03, n = 64) and OPL (12.54 ± 0.36 vs. 11.63 ± 0.21; p = 0.04, n = 64) in S1PR2 KO mice were significantly thicker than those in WT mice. These results suggest that S1PR2 may regulate the establishment of specific retinal layers, which is not different at 3 or 6 months of age and might affect overall retinal function.

### Role of S1PR2 in the expression of synaptic proteins

To further investigate the changes in the light responses, particularly the increased b-wave amplitude we observed in both the retinas of both dark and light-adapted S1PR2 KO mice, we examined the retinal synapses and neuronal morphologies in WT and S1PR2 KO mice using IHC assays and confocal fluorescence microscopy (**Table 1**).

Since we also observed an increase in OPL thickness by OCT, we wondered whether the increased a- and b-wave amplitude in S1PR2 KO mice resulted from an increased number of photoreceptors. Therefore, we quantitated the retinal cell densities and measured the expression of cell-specific markers. Confocal microscopy scans of DAPI-stained transverse retinal-cross sections from wild-type and S1PR2 KO mice demonstrates the preservation of all nuclear layers (**Figures 3A and 3D**). The G-coupled receptor protein rhodopsin found in rod cells regulates phototransduction upon light activation,[16] while cone arrestin is expressed in cone photoreceptors that contribute to high acuity color vision.[17] Thus, we evaluated rod and cone photoreceptors using antisera specific for rhodopsin (**Figures 3B and 3E**) and cone arrestin (**Figures 3C and 3F**), respectively. Fluorescence confocal microscopy showed that antibodies specific for rhodopsin clearly immunolabeled the rod outer segments but we observed no differences in signal between the retinas from WT and S1PR2 KO mice (**Figure 3H**). Similarly, fluorescence confocal microscopy with antibodies specific for cone arrestin allowed us to visualize the cone outer segments and terminals, yet we detected no differences in cone arrestin expression between the retinas of WT and S1PR2 KO mice (**Figure 3I**). These data demonstrate no changes in the number of photoreceptors (**Figure 3G**) or the expression pattern of rod and cone-specific proteins in the retinas of S1PR2 KO mice relative to those of the WT controls (**Figures 3H and 3I**).

**Figure 3.**
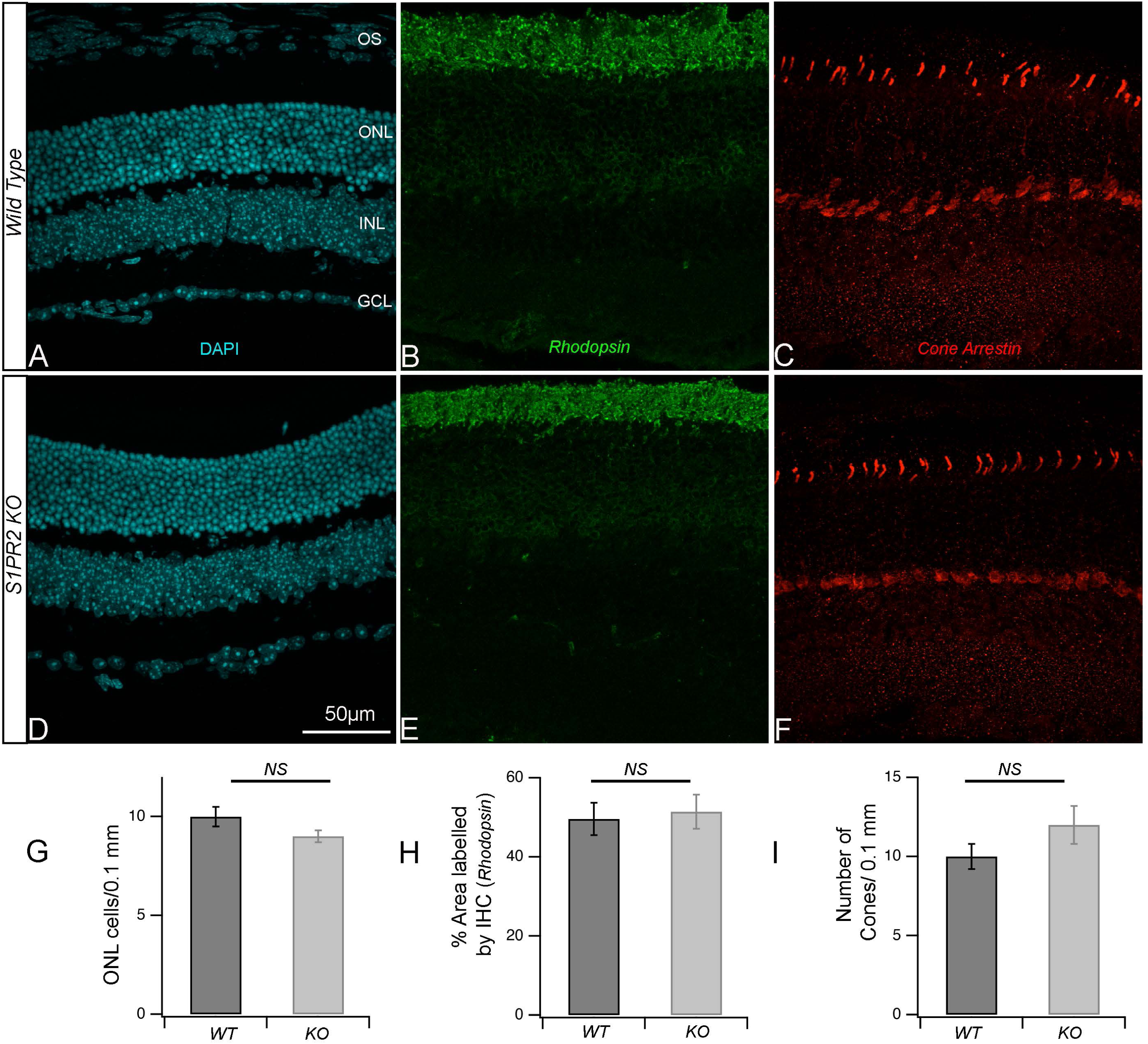
The retinas from S1PR2 KO mice retained WT cell densities in all nuclei layers and both rod and cone photoreceptors. (**A-F**) Transverse retinal cross-sections from 3-month-old WT control (**A-C**) or S1PR2 KO (**D-F**) mice were stained with DAPI (cyan, **panels A and D**) or immunostained with fluorescently labeled antibodies specific for the rod photoreceptor rhodopsin (green, **panels B and E**) or cone photoreceptor cone arestin (red, **panels C and F**). Scale bars: 50. Maximal intensity projections are shown. (**G-I**) Quantitative analyses for (**G**) as a function of the number of DAPI stained nuclei in the ONL per 0.1 mm area. Quantitative analyses of IHC for rhodopsin as a measure of the % of the total area that contained rods (**H**) or cone arrestin (**I**) as a measure of the number of cones. Abbreviations used: DAPI, 4’,6-diamidino-2-phenylindole; KO, knockout; NS, not significantly different; ONL, outer nuclear layer; OPL, outer plexiform layer; OS, outer segments; WT, wildtype. Data represent the average value of the area of expression in 4–6 animals sampled per condition in 3–5 independent experiments.

Next, we examined the cell densities of secondary order neurons in light-adapted transverse retinal-cross sections from WT and S1PR2 KO mice. In our previous study, we found S1PR2 expression in both photoreceptors and bipolar cells, and b-wave responses of light or dark-adapted retina are generated by both ON and OFF-type bipolar cells. Here we used fluorescently-labeled antibodies specific for the rod bipolar cell marker protein kinase C α (PKCα; **Figures 4A and 4D**) and cone bipolar marker secretagogin (**Figures 4B and 4E**). Quantitative analysis of the area labeled by PKCα and secretagogin antibodies revealed no differences between WT and S1PR2 KO mice (**Figures 4G and 4H**). In addition to bipolar cells forming synapses with photoreceptors, the dendrites of horizontal cells also connect with cone pedicles as lateral invaginating elements at the ribbon synapse, while the axonal processes connect to rod spherules in an equivalent synaptic configuration.[18] The immunoreactivity of the calcium-binding protein calbindin is most prominent in horizontal cells of the mouse retina.[19] Calbindin is widely distributed in retinal neuronal cells, where it protects cells against apoptosis.[20] Fluorescence confocal microscopy of retinas from WT and S1PR2 KO mice with antibodies specific for calbindin (**Figures 4C and 4F**) and quantitative analysis of the area labeled by calbindin antibodies (**Figure 4I**) revealed no differences in signal. Indeed RT-qPCR analysis further confirmed the preservation of retinal markers in S1PR2 KO mice (**Supplement Figure S2**). These data and DAPI labeling suggest that no gross changes in the neurons that comprise the OPL synapses in S1PR2 KO mice relative to those of WT mice.

**Figure 4.**
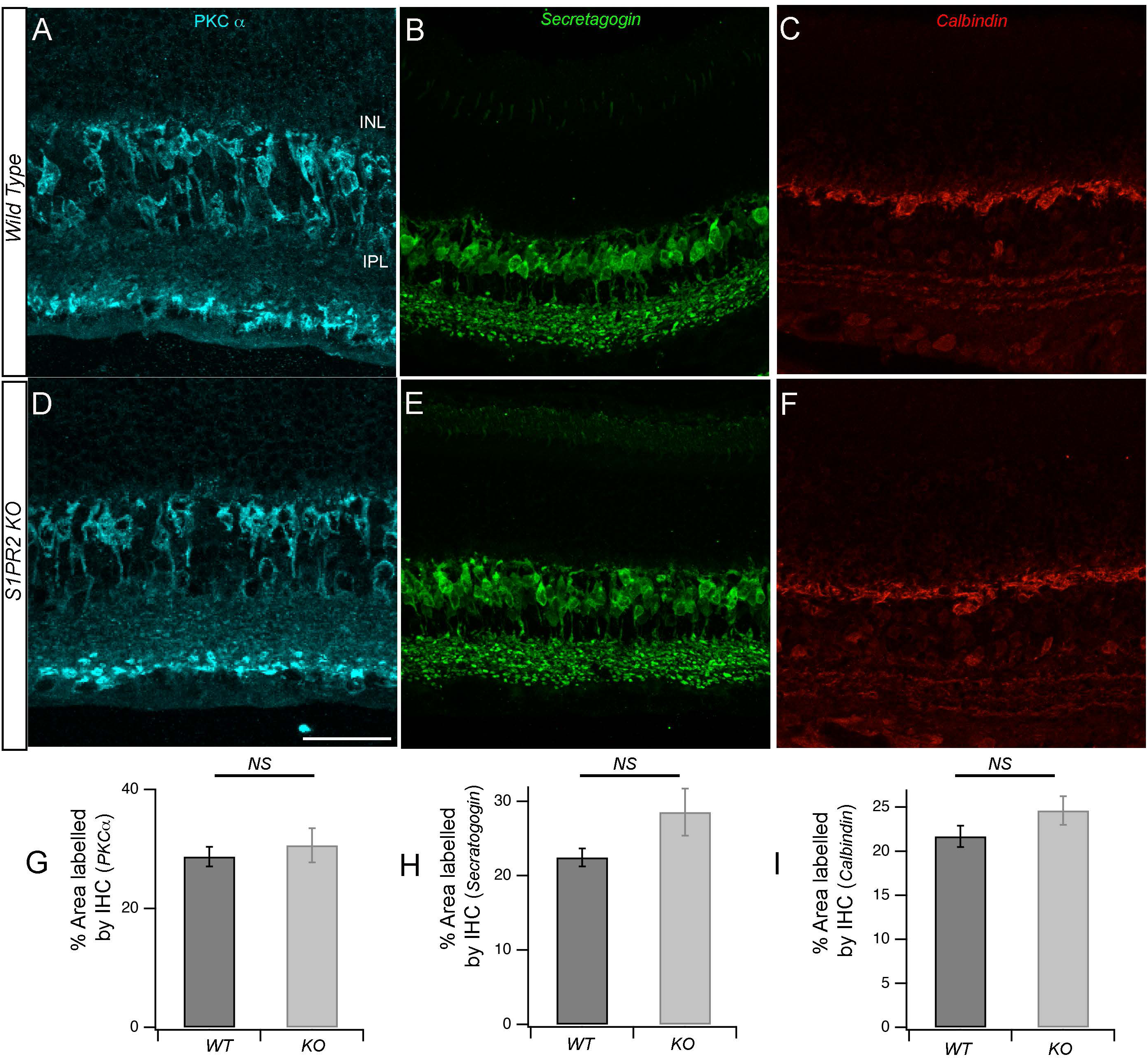
Secondary order neuron changes in S1PR2 KO mice. Immunostaining and quantitative analysis of rod bipolar cells (cyan, **A and D**), cone bipolar cells (green, **B and E**), and horizontal cells (red, **C and F**) from WT (**A-C**) or S1PR2 KO (**D-F**) mice. Scale bars: 50 Immunostaining with PKCα (**panels A and D**) showed no changes in the number of rod bipolar cells, as confirmed by the lack of statistically significant differences in the rod bipolar cell area of expression detected on quantitative analysis (**G**). Immunostaining with antibodies specific for secretagogin (**panels B and E**) or calbindin (**panels C and F**) revealed no changes in the number of cone bipolar cells or horizontal cells, respectively, as confirmed by the lack of statistically significant differences in the expression areas for secretagogin in cone bipolar cells (**H**) and calbindin in horizontal cells (**I**). Data represent the average value of the area of expression in 4–6 animals sampled per condition in 3–5 independent experiments. Abbreviations used: GCL, ganglion cell layer; KO, knockout; NS, not significantly different; ONL, outer nuclear layer; OPL, outer plexiform layer; WT, wild type.

Since photoreceptor and bipolar cell synapses at the OPL are the primary resources of the b-wave response and any changes during the establishment of these synapses can affect the overall visual pathway, we focused on the synaptic proteins that establish the OPL synapses. Second-order neurons in the OPL form specialized chemical ribbon synapses with photoreceptors neurons. Ribbon synapses possess specialized active zones that allow fast, precisely-timed signaling over prolonged periods. The main structural component of the synaptic ribbon is the ribeye protein. The unique N-terminus of ribeye is comprised of a proline-rich A-domain and a carboxyterminal B-domain that, except for its N-terminal 20 amino acids, is identical to the nuclear transcriptional co- repressor C-terminal binding protein 2 (CtBP2).[21, 22] We used antibodies specific for the ribeye *a domain* [23] to reveal the morphology and maturity of synaptic pre-synaptic protein (**Figures 5A and 5C**). Quantitative analysis showed that labeling of the ribeye a domain was stronger in the OPL of S1PR2 KO mice (27.4 ± 2.4%) than in age-matched WT mice (12.6 ± 1.5%; p<0.001), as shown in **Figure 5E**.

**Figure 5.**
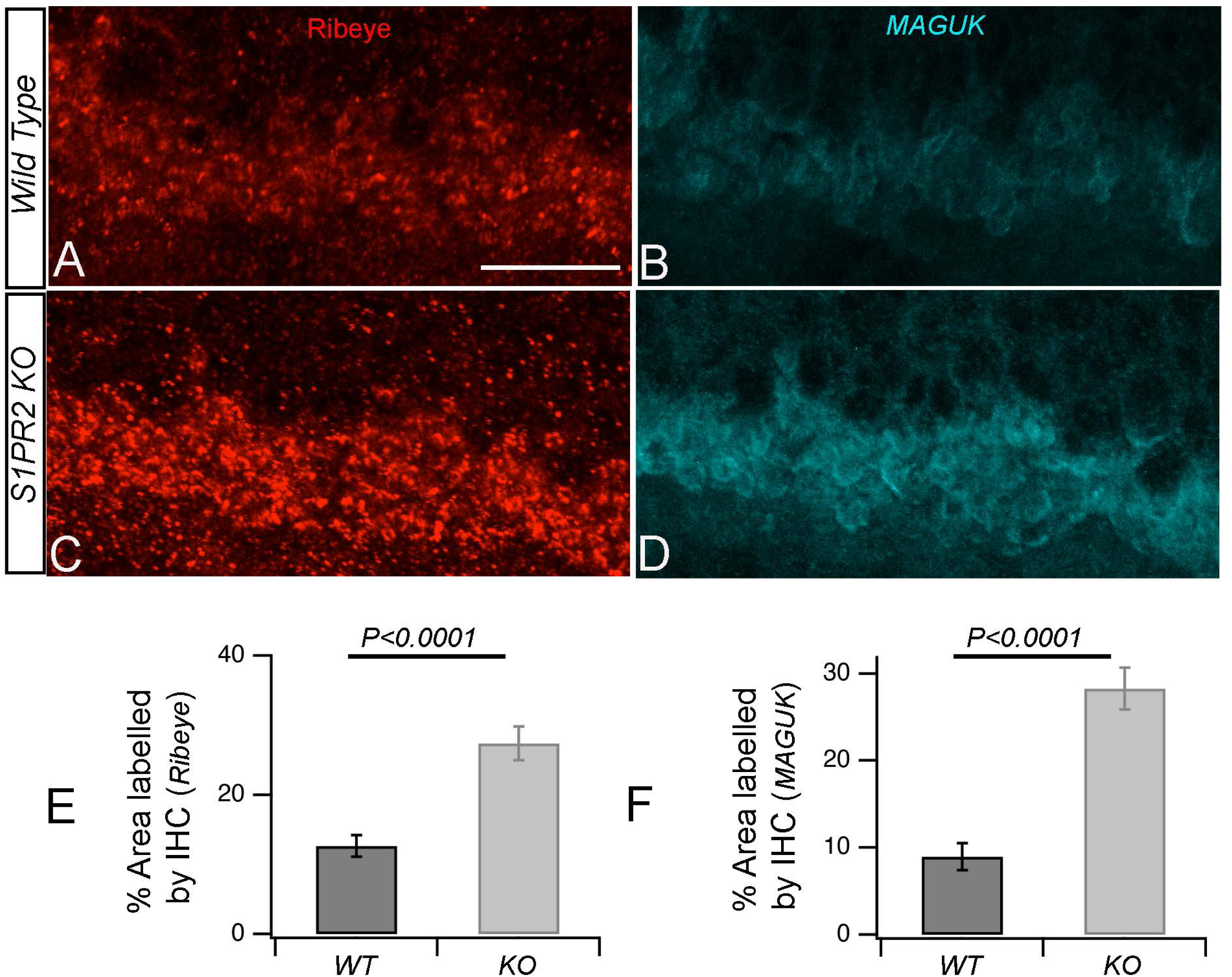
Synaptic structure in the OPL of WT and S1PR2 KO mice. (**A**) Transverse retinal cross-sections from WT (**A-B**) or S1PR2 KO (**C-D**) mice were immunostained with fluorescently labeled antibodies specific for synaptic ribbon marker ribeye (red, **A and C**) postsynaptic marker MAGUK (cyan, **B and D**). Maximal intensity projections are shown. Scale bars: 50 (**B**) Quantitative analyses of IHC for accumulation of ribeye and MAGUK proteins in the OPL represent the average value of the area of expression in 4–6 animals sampled per condition in 3–5 independent experiments. Statistical relevance was determined using a two-tailed *t*-test; *p* < 0.0001. Abbreviations used: IHC, immunohistochemistry; KO, knockout; OPL, outer plexiform layer; WT, wild type.

To determine whether this increased expression of the OPL presynaptic marker ribeye occurs simultaneously with the increase in its postsynaptic counterparts, we examined the expression pattern of a postsynaptic marker, namely the PSD-95 family membrane-associated guanylate kinase MAGUK, which assembles and regulates the postsynaptic densities of the excitatory synapse. [24] MAGUK regulates trafficking, targeting, and signaling of ion channels in neuronal synapses[25] and assembles and regulates the postsynaptic densities of the excitatory synapse. [24, 26] We immunolabeled transverse retinal-cross sections from WT and S1PR2 KO mice with antibodies specific for MAGUK and performed IHC analysis as before. We observed increased labeling of MAGUK in the retinal OPL of S1PR2 KO mice in a manner similar to that observed for the ribeye protein (**Figures 5B and 5D**). The quantitative analysis showed that the area labeled by the anti-MAGUK antibodies was 8.9 ± 1.5% for the retinas of WT mice, compared with 28.27 ± 2.4% for S1PR2 KO mice (*p*<0.001), as shown in **Figure 5F**.

We wondered whether the increased levels of ribeye protein in the retinas of S1PR2 KO mice were associated with increased synaptic vesicles to accommodate the aforementioned increased ERG b-wave response. The synaptic proteins synaptic vesicle glycoprotein 2 (SV2) are essential for pre- synaptic stabilization and for regulation of calcium and neurotransmitter release,[27] while synaptophysin contributes to synaptic vesicle formation and exocytosis and is essential for neurotransmitter delivery.[28] Interestingly, fluorescence confocal imaging with antibodies specific for SV2 (**Figures 6A and 6C**) and synaptophysin (**Figures 6B and 6D**) demonstrated increased immunolabeling of retinal sections from S1PR2 KO mice relative to those from WT mice. The quantitative analysis showed that the OPL area labeled by anti-SV2 antibodies was 16.7± 1.8% in the retinas from WT mice and 23.9 ± 2% in those from S1PR2 KO mice (p<0.05), as shown **Figure 6E**, and by anti-synaptophysin antibodies was 17.9± 2.5% in the retinas from WT mice and 31.1 ± 2% in those from S1PR2 KO mice (p<0.05), as shown in **Figure 6F**.

**Figure 6.**
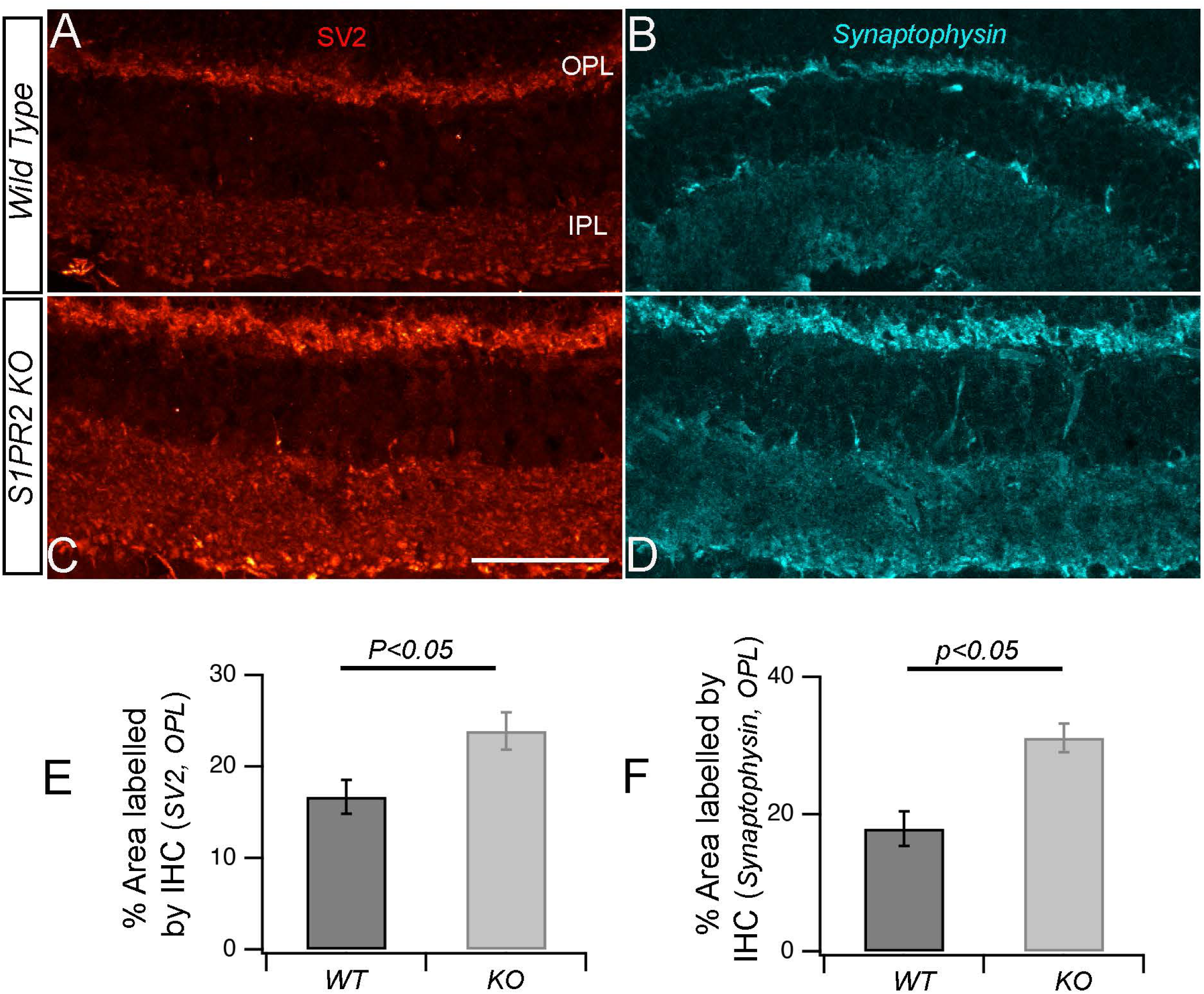
Synaptic vesicle protein accumulation in the OPL increased in SPR2 KO mice. (**A**) Transverse retinal sections from WT and S1PR2 KO mice were immunostained with fluorescently labeled antibodies for synaptic vesicle markers SV2 and synaptophysin. Maximal intensity projections are shown. Scale bar, 50 µm. Accumulation of SV2 and synaptophysin was increased significantly in S1PR2 KO mice relative to WT mice. (**B**) Quantitative analyses of IHC for SV2 and synaptophysin accumulation in WT and S1PR2 KO mice are shown by bar graphs. Data represent the average value of the area of expression in 6–9 animals sampled per condition in 3–5 independent experiments. Statistical relevance was determined using a two-tailed *t*-test; *p* < 0.05. Abbreviations used: IHC, immunohistochemistry; IPL, inner plexiform layer; KO, knockout; OPL, outer plexiform layer; SV2, synaptic vesicle protein 2.

### S1PR2 KO mice exhibit illumination-dependent changes in rod bipolar cell morphology and synaptic protein expression

In mammalian rod ON bipolar cells, PKCα expression is activity-dependent.[29, 30] We found that the dendrites of dark-adapted retinas were thicker, but those of light-adapted retinas were more branched in both WT and S1PR2 KO mice (compare **Figures 7A & 7B** with **Figures 7C & 7D**). The synaptic terminals of mammalian rod-ON bipolar cells and goldfish mixed rod–cone ON bipolar cells exhibit morphological changes in response to different light levels.[30] For example, in the light-adapted retina, the synaptic membranes in the axon terminals of rod bipolar cells were round with smooth convex curvature, while those in the dark-adapted retina exhibited irregular contours with numerous dimples and concave curvature.[30–32] We also observed similar changes in the terminal morphology of fluorescently-immunolabelled PKCα-rod bipolar cells from WT but not S1PR2 KO mice (compare **Figures 7E & 7F** with **Figures 7G & 7H**). As demonstrated in rat rod bipolar cells previously, [30] in light-adapted rod bipolar cells from WT mice, the immunoreactivity of PKCα were homogeneously distributed throughout the cytoplasm (**Figure 7E**), but in dark-adapted state, PKCα preferentially localized to the submembrane (**Figure 7F**). In contrast, PKCα expression in S1PR2 KO mice was largely homogeneous distribution throughout the cytoplasm in both light- and dark-adapted states (compare **Figures 7E & 7F** with **Figures 7G & 7H**).

**Figure 7.**
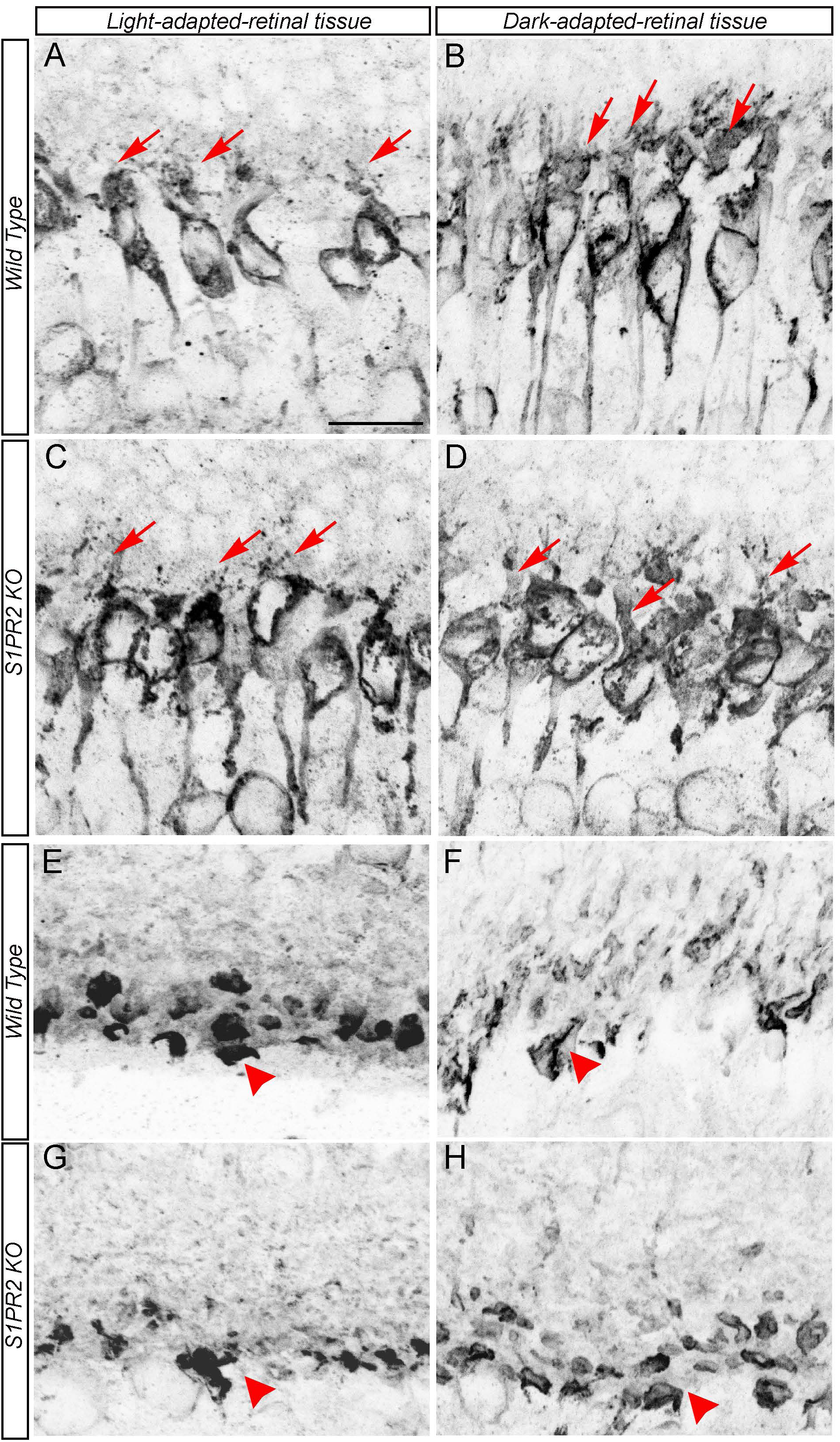
Rod bipolar cells from S1PR2 KO mice exhibited abnormal light-dependent protein kinase C alpha distribution relative to those in WT mice. Immunostaining of rod bipolar cell-specific marker PKCα in light-adapted (**left** panels **A, C, E, and G**) or dark-adapted (**right** panels **B, D, F, and H**) retinas from WT **(A-B, and E-F)** o4 S1PR2 KO mice **(C-D and G-H).** Rod bipolar cell somata exhibited light-dependent changes in dendritic arborization (**A-D,** see white arrows). Rod ON bipolar cells from WT mice exhibited light/dark-dependent morphological changes at the surface of their synaptic terminals **(E-F)**. In light-adapted retinas, axon terminals were round and PKCα immunoreactivity was homogeneously distributed **(E,** see red arrowheads), while in dark-adapted retinas, the terminals had irregular contours and PKCα immunoreactivity was localized in the submembrane compartment and staining of the more central cytoplasm was reduced in **(F,** see red arrowheads). However, such light/dark dependent PKCα expression was not evident in rod bipolar cell terminals of S1PR2 KO mice. Scale bars: 50 μm. Abbreviations used: KO, knockout; PKCα, protein kinase C alpha; WT, wildtype. Data represent the average value of the area of expression in 5–7 animals sampled per condition in 4–6 independent experiments.

To determine whether altered adaptation-dependent plasticity of rod bipolar cell axon terminal morphology was a general phenomenon in the S1PR2 KO mice, we examined the expression pattern of the illumination-dependent retinal protein ribeye[33–37] in rod bipolar cells from light- and dark-adapted WT and S1PR2 KO mice. Fluorescence confocal analysis with antibodies specific for ribeye showed that the OPL of dark-adapted retinas was more heavily stained than light-adapted retinas in both WT and S1PR2 KO mice, but in both light and dark-adapted retinas, both the number and intensity of immunoreactive ribeye puncta were higher in S1PR2 KO mice than in WT mice (**Figures 8A-D and I**). We also found that the postsynaptic marker MAGUK follows an illumination-dependent expression pattern similar to that of ribeye in the retinas of both WT and S1PR2 KO mice, with higher expression in the light and dark-adapted retinas of S1PR2 KO mice than in those of WT mice (**Figures 8E-H and J**).

**Figure 8.**
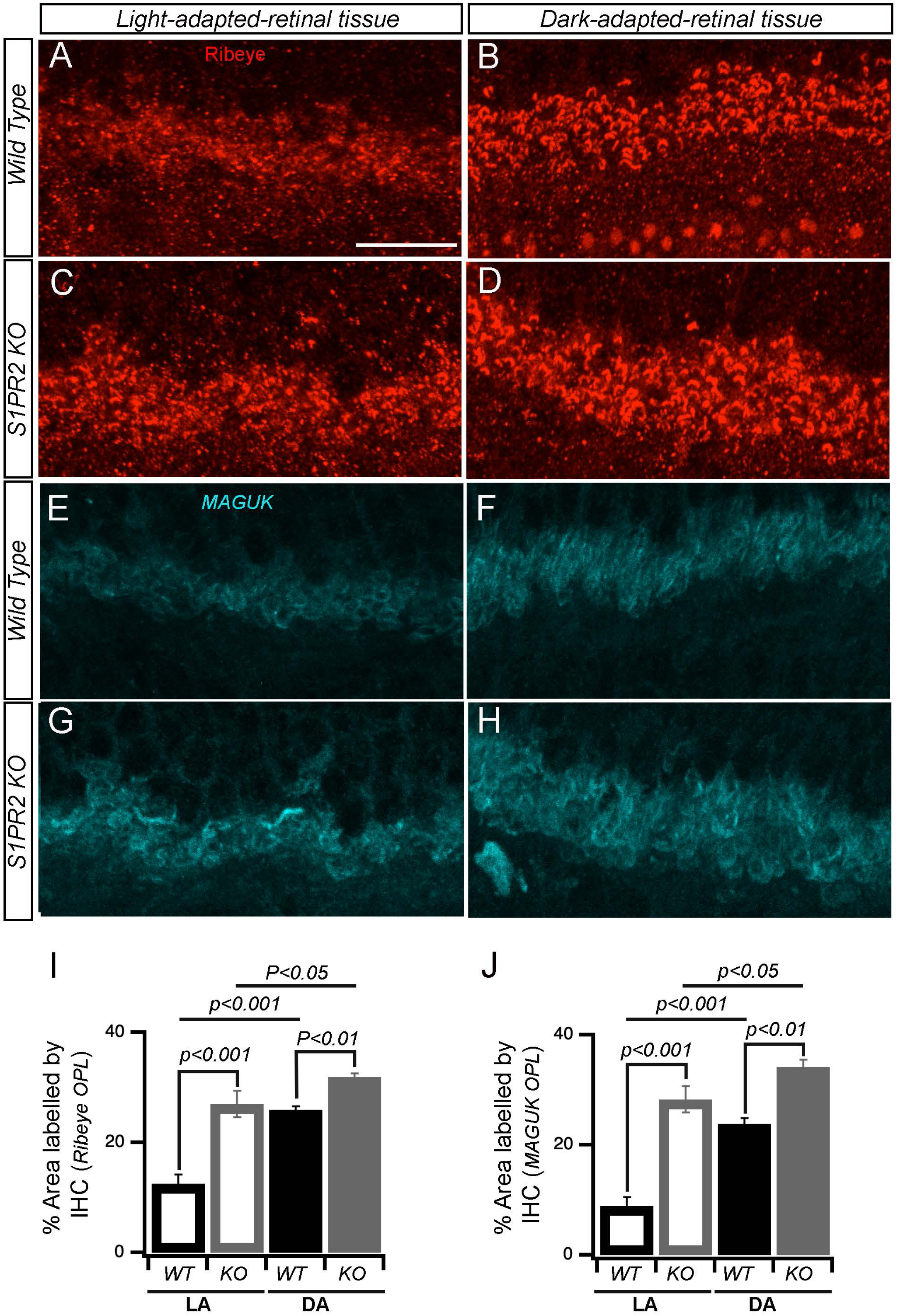
S1PR2 KO mice exhibited diminished illumination-dependent remodeling of the photoreceptor active zone. Immunostaining of photoreceptor synapses from light-and dark-adapted retinas immunolabelled with polyclonal antibodies against the ribeye protein. Representative confocal images of dark-and light-adapted retinas from WT (**A-B and E-F**) and S1PR2 KO mice (**C-D and G-H**) are shown. Ribeye immunosignals were more intense and a greater number of immunoreactive ribeye puncta were observed in light- and dark-adapted retinas from S1PR2 KO mice (**C-D**) than in those from WT mice (**A-B**). Expression of the post- synaptic marker MAGUK demonstrated a similar pattern in light-and dark-adapted retinas from WT (**E-F**) and S1PR2 KO mice (**G-H**), which higher expression in retinas at both light levels from S1PR2 KO mice than those from WT mice. Scale bar: 50 µm. Abbreviations used: KO, knockout; WT, wildtype. (**I-J**) Quantitative analyses of IHC for accumulation of ribeye and MAGUK proteins in the OPL in LA and DA state represent the average value of the area of expression in 4–6 animals sampled per condition in 3–5 independent experiments. Statistical relevance was determined using a two-tailed *t*-test;. Abbreviations used: IHC, immunohistochemistry; KO, knockout; OPL, outer plexiform layer; WT, wild type; LA, light-adapted; DA, dark-adapted.

To determine the effects of light-dependent changes in the number of OPL synapses, we co-immunolabeled the retinas of WT and S1PR2 KO mice with antibodies specific for the presynaptic marker ribeye and the postsynaptic markers PKCα (**Figures 9A-D**) or MAGUK (**Figures 8E-H**) and performed IHC analysis as before. We observed that the labeling of ribeye partially overlapped that of the dendritic tips of the bipolar cells stained for PKCα or MAGUK in both WT and S1PR2 KO mice. Though the overlap of pre-and post-synaptic markers in the OPL followed similar patterns in WT and S1PR2 KO mice, the number of fluorescently labeled cells with ribeye and PKCα (**Figures 9I)** or MAGUK (**Figures 9J)** was greater in the retinas of S1PR2 KO mice than in those of WT mice at both light levels (compare **Figures 9A & 9B** with **Figures 9B & 9D and Figures 9E & 9F** with **Figures 9G & 9H**).

**Figure 9.**
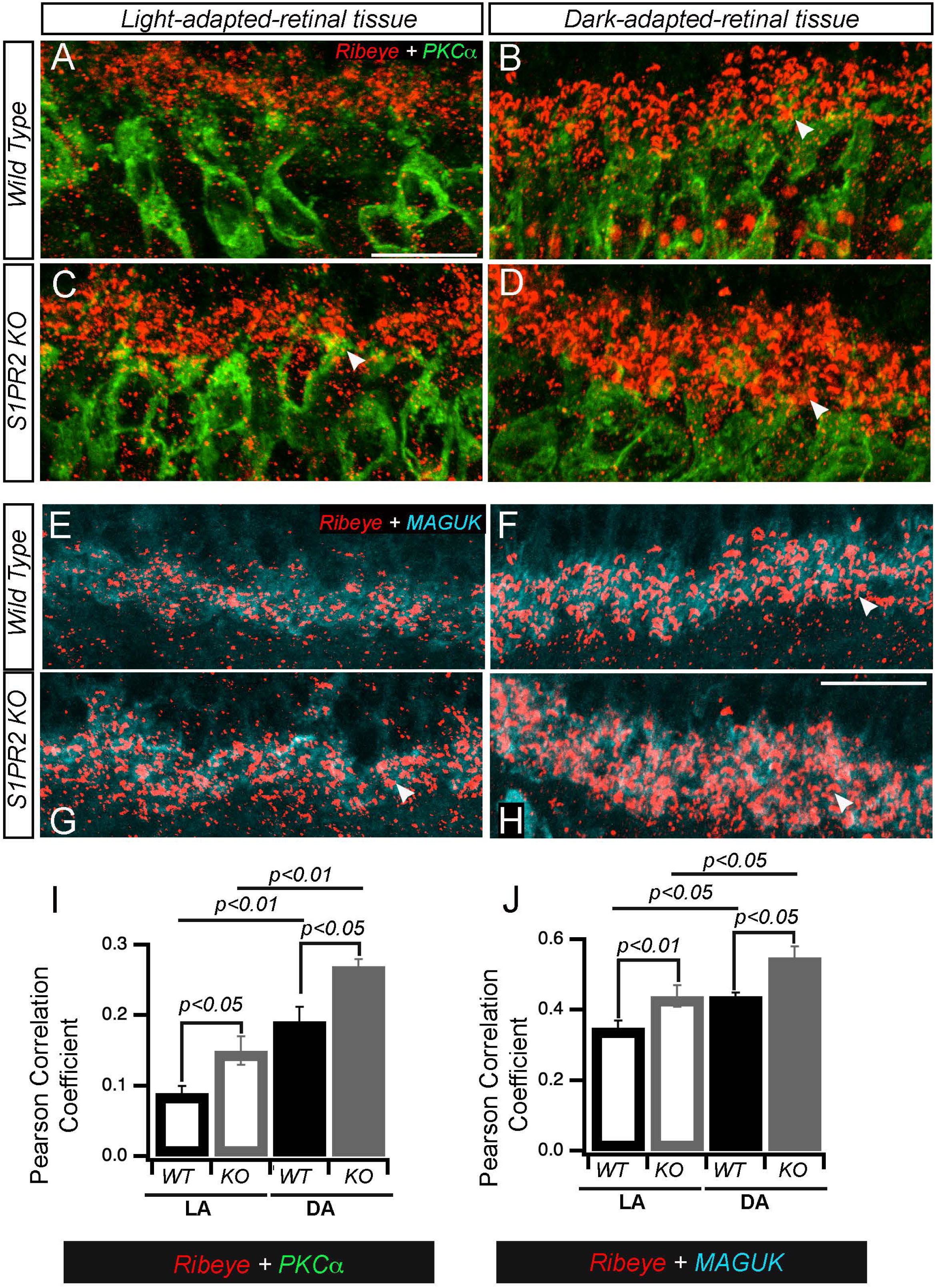
The retinas of S1PR2 KO mice exhibited altered photoreceptor ribbon synapses with postsynaptic densities (PSDs) in INL dendrites. (**A-D**)Transverse cross-sections of light- adapted (**left** panels **A, C, E, and G**) or dark-adapted (**right** panels **B, D, F, and H**) retinas from WT (**A-B and E-F**) or S1PR2 KO mice (**C-D and G-H**) were double immunostained with fluorescently labeled antibodies specific for either (**A-D**) the ribbon-specific marker ribeye (red) and the rod bipolar cell-specific marker PKCα (green) or (**E-H**) ribeye (red) and the postsynaptic marker MAGUK (cyan). Maximal intensity projections are shown; arrowheads indicate potential areas of colocalization. Scale bar, 50 µm. Abbreviations used: KO, knockout; MAGUK, membrane-associated guanylate kinase; PKCα, protein kinase C alpha; PSD, postsynaptic densities; WT, wildtype. (**I-J**) Pearson correlation analyses of IHC for accumulation of PKCα and ribeye or ribeye and MAGUK proteins in the OPL in LA and DA state represent the average value of the area of expression in 4–6 animals sampled per condition in 3–5 independent experiments. Statistical relevance was determined using a two-tailed *t*-test; Abbreviations used: IHC, immunohistochemistry; KO, knockout; OPL, outer plexiform layer; WT, wild type; LA, light-adapted; DA, dark-adapted.

### The role of S1PR2 in the expression of genes encoding synaptic proteins

To test whether the expression of synaptic proteins also differed *in vivo*, we isolated total RNA from the retinas of WT and S1PR2 KO mice and performed RT-qPCR analysis. We observed significantly greater expression of the pre-synaptic proteins bassoon and ribeye and the post- synaptic protein metabotropic glutamate receptor 6 (mGluR6) in S1PR2 KO mice than in WT mice (**Figure 10**). Bassoon proteins regulate clustering of pre-synaptic vesicles,[38] ribeye allows tethering of synaptic vesicles near the active zone membrane in ribbon synapses,[22] and mGluR6 is the primary post-synaptic glutamate receptor of retinal ON bipolar cells that receive glutamatergic input from photoreceptors.[39] The expression of several other pre- and post- synaptic proteins exhibited an upward trend that was not statistically significant (**Figure 10**). This data suggests that S1PR2 may play a role in the regulation of expression for synaptic genes, leading to the increased synaptic protein expression and b-wave amplitudes we observed in this study.

**Figure 10.**
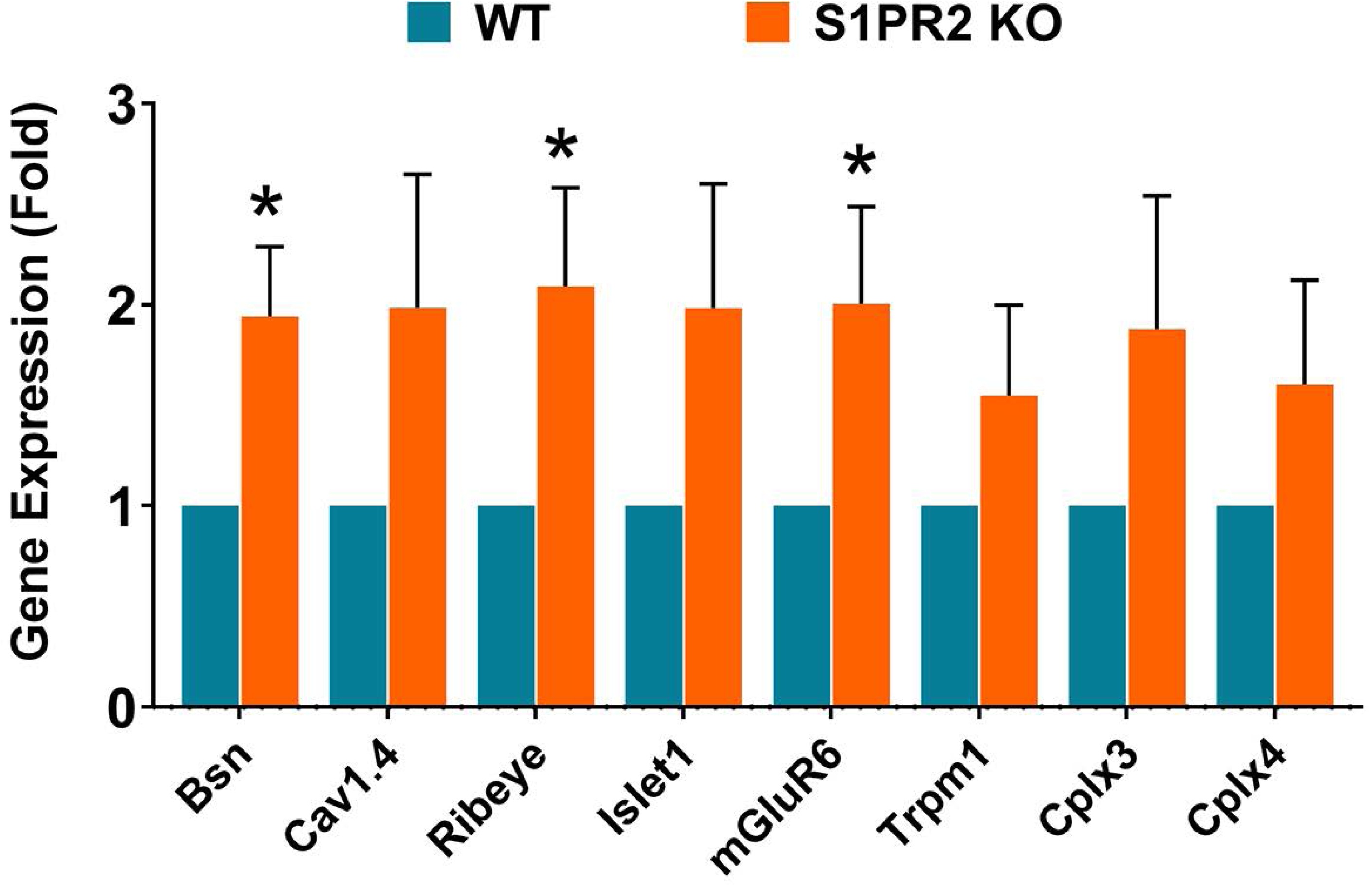
Expression of retinal pre- and post-synaptic markers is significantly increased in S1PR2 KO mice relative to WT mice. Statistical analysis of gene expression of pre- and post- synaptic markers in the retinas of WT (blue) and S1PR2 KO (orange) mice by RT-qPCR. Retinal expression of bsn, ribeye, and mGluR6 was significantly elevated in S1PR2 KO mice relative to that in WT mice. Shown is mean fold over control ± SEM. \**p* ≤ 0.05 indicates a significant difference between groups of WT and S1PR2 KO mice; N = 6. Abbreviations used: Bsn, bassoon; Cav1.4, voltage-dependent calcium channel subtype 1.4; Cplx3 and Cplx4, complexins 3 and 4; KO, knockout; mGluR6, metabotropic glutamate receptor 6; Trpm1, transient receptor potential cation channel M1; WT, wildtype.

## Discussion

To the best of our knowledge, this is the first investigation to compare the roles of S1PR2 in the neural retina of WT and S1PR2 KO mice. We found that S1PR2 affected retinal layer thickness, expression of synaptic genes and proteins, and light responses at the functional and cellular level and thus played an active role in synaptic structure and retinal function.

To reveal the roles of S1PR2 in the rod and cone pathways, we obtained ffERG measurements in the scotopic state for the rod pathway and photopic measurements for the cone pathway in response to brief light flashes of different intensities. The ffERG measures mass responses of the retina from several sources, which depends on both strength of the stimulus flash and the state of adaptation.[40] The output of this type of ERG waveform has two important components, the a-wave, and the b-wave. The a-wave is a short negative wave potential that indicates the activity of rod and cone photoreceptors in the outer retina under the scotopic and photopic state.[41] The a-wave is followed by the b-wave, a positive wave potential that reflects the activity of cells in the inner nuclear layer, especially synaptic transmission between photoreceptors and bipolar cells in the inner layers of the retina.[42] Under scotopic conditions, S1PR2 KO mice exhibited significantly higher a-amplitudes than WT mice, which reflects the increased activity of the photoreceptors upon stimulation. Under these same conditions, S1PR2 KO mice also exhibited significantly higher b-wave amplitudes than WT mice, which indicates the hyperactivity of synaptic transmission in the OPL in S1PR2 KO mice. The expression of S1PR2 in the photoreceptors and bipolar cells of mouse retina, as we reported previously, and the lower ERG amplitude of a- and b-waves in WT mice relative to S1PR2 KO mice suggests that S1PR2 may have an inhibitory role in establishing light responses.

We showed that S1PR2 did not regulate ONL cell density, which therefore did not contribute to the increased light responses we observed in S1PR2 KO mice relative to WT controls. Instead, the results of our differential labeling of pre-and post-synaptic markers in the OPL in WT and S1PR2 KO mice indicated an important role of S1PR2 in regulating synaptic neurotransmission. We observed enhanced expression of four different synaptic proteins in S1PR2 KO mice, namely ribeye, MAGUK, synaptic vesicle protein 2 (SV2), and synaptophysin. Ribeye is a ribbon-specific pre-synaptic protein found in the retina, inner ear, and pineal gland that tethers vesicles near the active zone.[22] Its increased expression in S1PR2 KO mice suggests that S1PR2 might regulate the size or the number of ribbon synapses. We also observed changes in the expression of the MAGUK postsynaptic protein in the OPL of S1PR2 KO mice relative to that of WT mice. MAGUK proteins are involved in the trafficking, signaling, and positioning of ion channels at specific cellular sites[25] and MAGUK p55 scaffold protein 4 (MPP4) is expressed in bipolar cell dendrites and their axonal terminals.[43] Thus, we hypothesize that the hyperactivity of ERG waveforms we observed in S1PR2 KO mice is likely due to the enhancement of ribeye and MAGUK expression in the pre-and post-synapse and the strengthening of OPL excitatory synapses.

We also observed the upregulation of two synaptic vesicle proteins, SV2, and synaptophysin. SV2 regulates the release of calcium and neurotransmitters,[27] synaptophysin contributes to synaptic vesicle formation and exocytosis.[28] The enhancement of synaptic vesicle pools and that the ribeye protein affects the tethering of synaptic vesicles to the ribbon further validates our hypothesis that the enhanced b-wave in S1PR2 KO mice is likely due to the strengthening of OPL excitatory synapses and that the increase in ERG b-wave amplitude in S1PR2 KO mice is likely due to enhanced synaptic transmission in the OPL. Further enhancement of the overlap of pre-and post-synaptic markers in the OPL in S1PR2 KO mice relative to that observed in WT mice at both light levels we examined (**Figure 9**) may suggest the underlying mechanisms by which scotopic and photopic b-wave responses increased in S1PR2 KO mice.

The increased synaptic activity of S1PR2 KO mice was supported by the results of our RT-qPCR analysis, which revealed upregulation in S1PR2 KO mice of three genes encoding synaptic proteins that are crucial for synaptic transmission, namely bassoon, ribeye, and mGluR6. Presynaptic OPL active zone proteins ribeye and bassoon[38] organize calcium channels and vesicles at release sites,[44] while mGluR6 is the main post-synaptic glutamate receptor of retinal ON bipolar cells, which receive glutamatergic input from photoreceptors.[39] Collectively, the results of RT-qPCR and immunoassays suggest that S1PR2 is involved in the regulation of synaptic protein expression. Thus, S1PR2 KO may lead to increased expression of synaptic markers, further validating our hypothesis of strengthening OPL synapses as a possible explanation for the increase in a- and b-wave amplitudes.

The retina exhibits numerous biochemical, physiological, structural, and synaptic morphological changes in activity-related dynamics and transition from light to darkness. The fact that our data show that the classic changes in PKCα expression in mammalian rod ON bipolar cells were absent in S1PR2 KO mice raises the possibility that S1PR2 signaling could involve circadian rhythms. The Nogo-A signaling pathway that is involved in regulating the visual system plays an important role in different neurodegenerative and retinal diseases [45] and is critical for regulating the circadian rhythms in other tissues.[46] Nogo-A signaling strengthens inhibitory synaptic transmission by regulating GABA receptors in pyramidal hippocampal neurons.[47, 48] Interestingly, the inhibition of Nogo-A signaling is mediated by S1PR2, which positively regulates the calcium-dependent clustering and localization of GABAA receptors at synapses.[1] In previous studies, inhibition of GABA function or elimination of GABAC expression resulted in a higher baseline level of a- and b-waves.[49]. Therefore, it is possible that S1PR2 regulates the inhibitory/excitatory balance reflected in the a- and b-wave amplitudes via a GABAergic mechanism within the Nogo-A signaling pathway. By the same token, increased expression of ribeye in S1PR2 KO mice suggests that S1PR2 may regulate the size or the number of ribbon synapses, which could occur through the loss of inhibitory GABA signal transmission between horizontal cells and photoreceptors.[50] Future studies will be needed to explore the expression of circadian clock genes, GABA, and Nogo-A signaling in S1PR2 KO mice to further understand the mechanisms underlying the altered synaptic expression and light responses we observed in S1PR2 KO mice during the present study.

## Conclusions

Overall, our studies provide the first evidence for the role of S1PR2 in light response, retinal structure, and synaptic transmission. S1PR2 also modulates retinal responses to light/dark adaptation, and in this study, S1PR2 interfered with the expression of important pre- and post- synaptic genes and proteins. S1PR2 signaling is likely to play a key role in balancing excitatory-inhibitory transmission, albeit future investigation will be required to uncover its underlying mechanism. One possibility is that S1PR2 may facilitate the inhibition of GABAergic synapses via Nogo-A signaling.[1] Together, these results suggest that S1PR2 plays an important role in the light responses of the retina and provides a useful target for the future development of novel therapeutics to treat and/or prevent different retinal diseases. These results allow future research to unveil the underlying mechanisms of S1PR2 in the neural retina to understand better the observed interactions and their role in neuronal diseases, malfunctioning, development, and aging.

## Supporting information

Supplmental Figure-1_3 mo ERG

Supplmental Figure-2_ Retinal Cell Markers Figure

## Author Declarations

### Ethics approval and consent to participate

Not applicable

### Author Consent

All the authors have seen the manuscript and approved it for publication.

### Availability of data and materials

All materials and data will be available from the corresponding author following the University of Tennessee’s policy of sharing research materials and data.

### Competing interests

None

## Funding

1. National Eye Institute of the National Institutes of Health under Award Number R01EY030863 (TV)
2. UTHSC College of Medicine Faculty Research Growth Award (TV)
3. National Eye Institute grants [EY022071, R01 EY031316] (NM)
4. US Department of Defense office of the Congressionally Directed Medical Research Programs (CDMRP) Vision Research Program grant [W81XWH-20-1-0900] (NM)
5. US Veterans’ Administration BLRD Merit Award [I01BX004893] (NM)
6. Research to Prevent Blindness Inc., USA (NM)
7. Intramural Research Program of the National Institutes of Health, National Institute of Diabetes and Digestive and Kidney Diseases to R.L.P.

## Author Contributions

Abhishek P Shrestha: performed experiments, prepared illustrations, and edited the manuscript; Megan Stiles: performed experiments and edited the manuscript; Richard C. Grambergs: performed experiments, analyzed data, and edited the manuscript; Johane M. Boff: analyzed data, prepared illustrations, wrote the first draft and edited the manuscript; Saivikram Madireddy: analyzed data, prepared illustrations, and edited the manuscript; Koushik Mondal: performed experiment and edited the manuscript; Rhea Rajmanna: analyzed data, prepared illustrations, and edited the manuscript; Hunter Porter: performed experiment, edit the manuscript; David Sherry: performed experiment, edited the manuscript; Richard L. Proia: provided resources and edited the manuscript; Thirumalini Vaithianathan: Designed experiment, obtained IACUC approvals, provided the facility, analyzed data, wrote the first draft, and edited manuscript; Nawajes Mandal: Designed experiment, obtained IACUC approvals, provided the facility, analyzed data, and edited manuscript.

## Acknowledgments

Research reported in this publication was supported by the National Eye Institute of the National Institutes of Health under Award Number R01EY030863 and UTHSC College of Medicine Faculty Research Growth Award to T.V. and National Eye Institute grants [EY022071, R01 EY031316], US Department of Defense office of the Congressionally Directed Medical Research Programs (CDMRP), Vision Research Program grant [W81XWH-20-1-0900], US Veterans’ Administration BLRD Merit Award [I01BX004893], and Research to Prevent Blindness Inc., USA to M.N, and by the Intramural Research Program of the National Institutes of Health, National Institute of Diabetes and Digestive and Kidney Diseases to R.L.P.

The authors would like to thank Dr. Kyle Johnson Moore in the UTHSC Office of Scientific Writing for the critical review and editing of the manuscript.

## Disclosure of potential conflicts of interest

None

## Research involving Human Participants

Not applicable.

## Animals

Animal studies approved by the UTHSC IACUC committee. Approval # 19-0104

---

**Supplement Figure S1. S1PR2 knockout (KO) mice exhibited higher baseline ERG responses than did wild-type (WT) at 3 months of age.** Shown are statistical analysis of scotopic ERG A- and B-wave amplitudes 3-month-old WT (blue) and S1PR2 KO (orange) mice. At higher flash intensities (400 and 2000 cd.s/m^2^), S1PR2 KO mice demonstrated significantly higher mean scotopic A-wave (A) and B-wave (B) ERG responses than WT mice. Shown are mean values ± SEM. \**p* < 0.05 indicate significant differences between groups of WT and S1PR2 KO mice (N = 4 mice per genotype). Abbreviations used: ERG, electroretinography; KO, knockout; WT, wildtype.

**Supplement Figure S2. Expression of retinal markers in WT and S1PR2 KO mice.** Statistical analysis of gene expression of retinal markers in the retinas of WT (blue) and S1PR2 KO (orange) mice by RT-qPCR. Retinal expression of CD68/LAMP4, Opn1mw, Opn1sw, PKCα, syntaxin 1A, Thy1/CD90, rod arrestin, and rhodopsin did not significantly differ between WT and S1PR2 KO mice. Shown is the mean fold value over control ± SEM (N= 6). Abbreviations used: CD68/LAMP4, cluster of differentiation 68/lysosomal-associated membrane protein 4; KO, knockout; Opn1mw, medium wave-sensitive opsin 1; Opn1sw, short wave-sensitive opsin 1; PKCα, protein kinase C alpha; Trpm1, Thy1/CD90, Thy1 cell surface antigen/cluster of differentiation 90; WT, wildtype.

## Notes

### Competing Interest Statement

The authors have declared no competing interest.

