## Supplementary figures and images for "The role of sphingosine-1-phosphate receptor 2 in mouse retina light responses"

### Supplmental Figure-1_3 mo ERG.tif

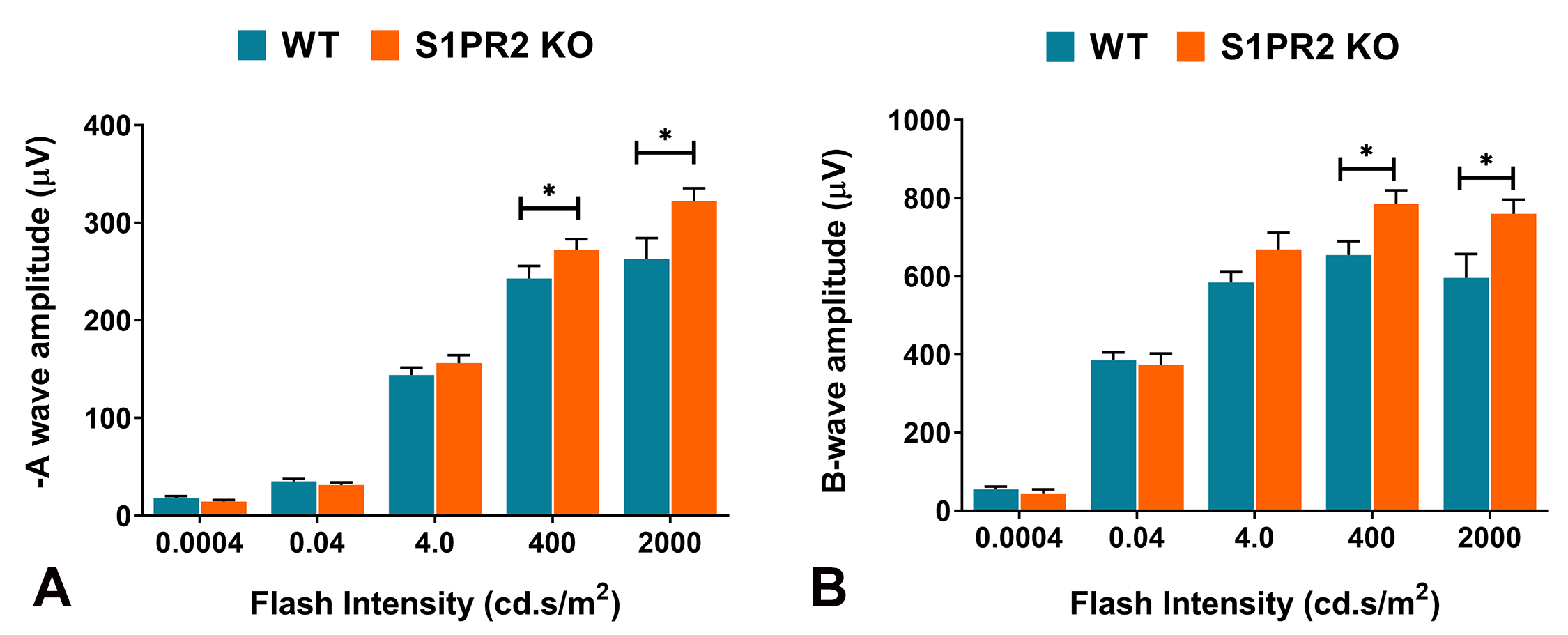

### Supplmental Figure-2_ Retinal Cell Markers Figure.jpg

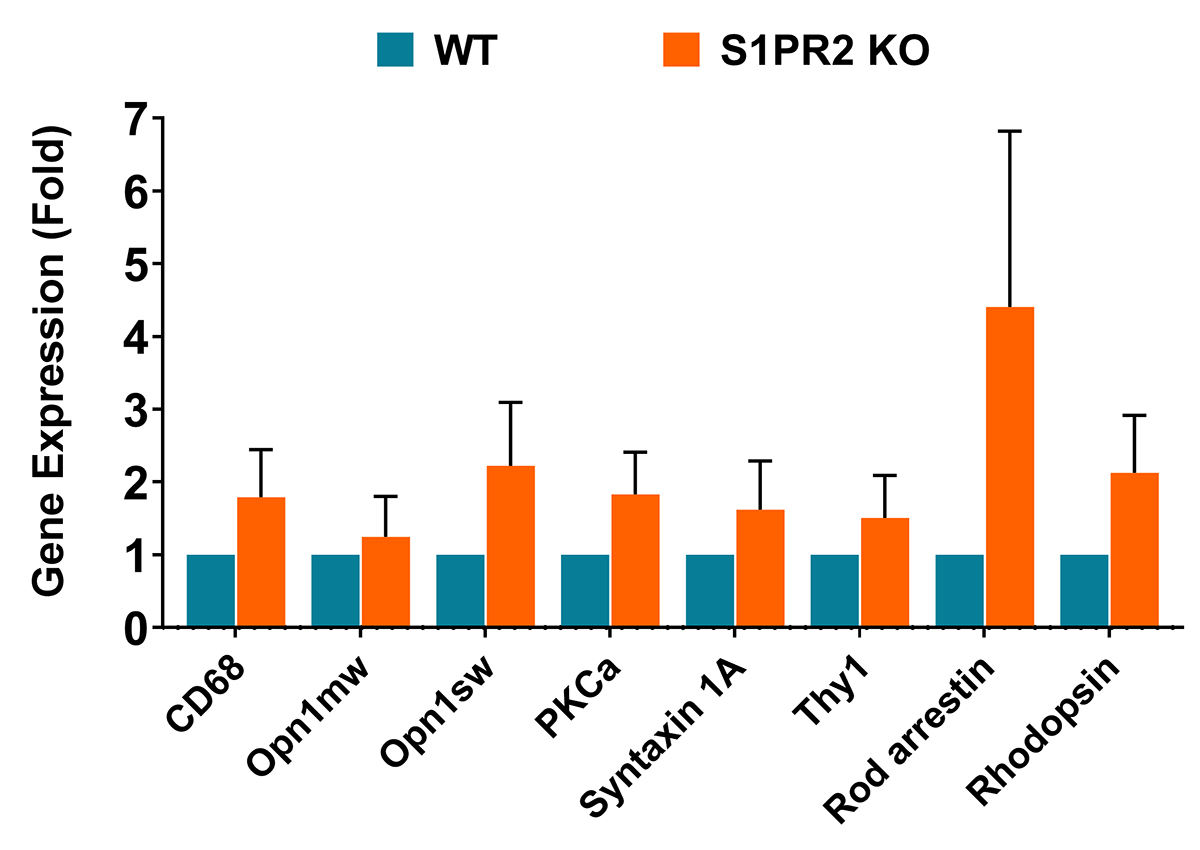
